# Biochemical reassessment of the activation of PANK1 by SGLT2 inhibitors

**DOI:** 10.64898/2026.11.09.765291

**Authors:** Zihao Guo, Jiahe Zang, Lei Chen

## Abstract

Although SGLT2 inhibitors (SGLT2i) show cardiovascular benefits in addition to glycemic control, their molecular target for cardiovascular effect remains unclear. A recent study reported that two SGLT2i, empagliflozin and dapagliflozin, directly activate pantothenate kinase 1 (PANK1), which mediate their cardioprotective effect. PANK1 phosphorylates pantothenate with ATP to yield 4′-phosphopantothenate and ADP, representing the rate-limiting enzyme of coenzyme A (CoA) biosynthesis. We therefore reassessed this claim using purified recombinant human PANK1. The malachite green assay detected free inorganic phosphate (Pi) but neither ADP nor 4′-phosphopantothenate and its signals do not reflect PANK1 activity. In the ADP-Glo assay, empagliflozin and dapagliflozin did not stimulate PANK1 under either basal conditions or acetyl-CoA feedback inhibition, while the activator PZ-2891 enhanced the activity of PANK1. These results show that SGLT2i do not directly activate PANK1, challenging the proposed PANK1-dependent cardioprotection mechanism.

## 1. Introduction

SGLT2 is mainly expressed on the apical membrane of epithelial cells in the renal proximal tubule and mediates the reabsorption of approximately 90% of the glucose filtered by the glomerulus, thereby playing an important role in whole-body glucose homeostasis^1^. SGLT2i, including empagliflozin and dapagliflozin, are small-molecule compounds derived from the natural product phlorizin^2^. These agents share a common modular structure consisting of a β-D-glucopyranose core, a proximal aromatic ring, and a distal hydrophobic ring^2^. SGLT2i occupies the sugar substrate binding site and the extracellular vestibule of SGLT2 and locks the transporter in the outward-facing conformation, arresting the alternating-access cycle required for Na□-coupled glucose transport across the membrane^3,4^. Consequently, SGLT2i block glucose reabsorption in kidney, induce pronounced glucosuria, and lower blood glucose independently of insulin secretion^5^. Therefore, SGLT2i are widely prescribed for the treatment of T2DM ^5^.

Unexpectedly, large-scale randomized clinical trials demonstrated that SGLT2i show benefits extending well beyond glycemic control^6^. In patients with T2DM and established atherosclerotic cardiovascular disease, empagliflozin (EMPA-REG OUTCOME) ^7^ and canagliflozin (CANVAS)^8^ reduced major adverse cardiovascular events, cardiovascular death, and hospitalization for heart failure, whereas dapagliflozin (DECLARE-TIMI 58)^9^ consistently lowered heart-failure hospitalization. Subsequent heart-failure-specific trials, DAPA-HF^10^ and EMPEROR-Reduced ^11^for HFrEF, and DELIVER^12^ and EMPEROR-Preserved^13^ for HFpEF, further showed that SGLT2i reduce heart-failure hospitalization and cardiovascular death for both HFrEF and HFpEF. Despite these robust and reproducible cardiovascular benefits, the molecular mechanisms underlying this phenomenon remain incompletely understood.

A recent study reported that SGLT2i, including empagliflozin and dapagliflozin, directly activate human cardiac PANK1 and proposed this mediates the cardiovascular benefits of these drugs^14^. PANK1 catalyzes the ATP-dependent phosphorylation of pantothenate to yield 4′-phosphopantothenate and ADP (Fig. 1A, B)^15^. This reaction is the first committed and rate-limiting step in CoA biosynthesis^15^. PANK1 activity is subject to negative feedback regulation by the CoA species: free CoA and CoA thioesters, including acetyl-CoA, malonyl-CoA, and long-chain acyl-CoA, which bind to PANK1 and inhibit its activity (Fig. 1C)^15^. PANK1 functions as a homodimer. The well-characterized PANK activator PZ-2891 occupies the pantothenate substrate binding site (Fig. 1D)^16^. At substoichiometric occupancy, binding of PZ-2891 to one protomer locks the other protomer in a catalytically active conformation that is insensitive to inhibition by endogenous CoA species, thereby activating the enzyme. However, at saturating concentrations, PZ-2891 binds both pantothenate pockets and acts as an orthosteric inhibitor (Fig. 1D)^16^. Here, we investigated whether SGLT2i can directly activate purified PANK1 using an assay that reports the PANK1-catalyzed reaction.

**Fig. 1.**
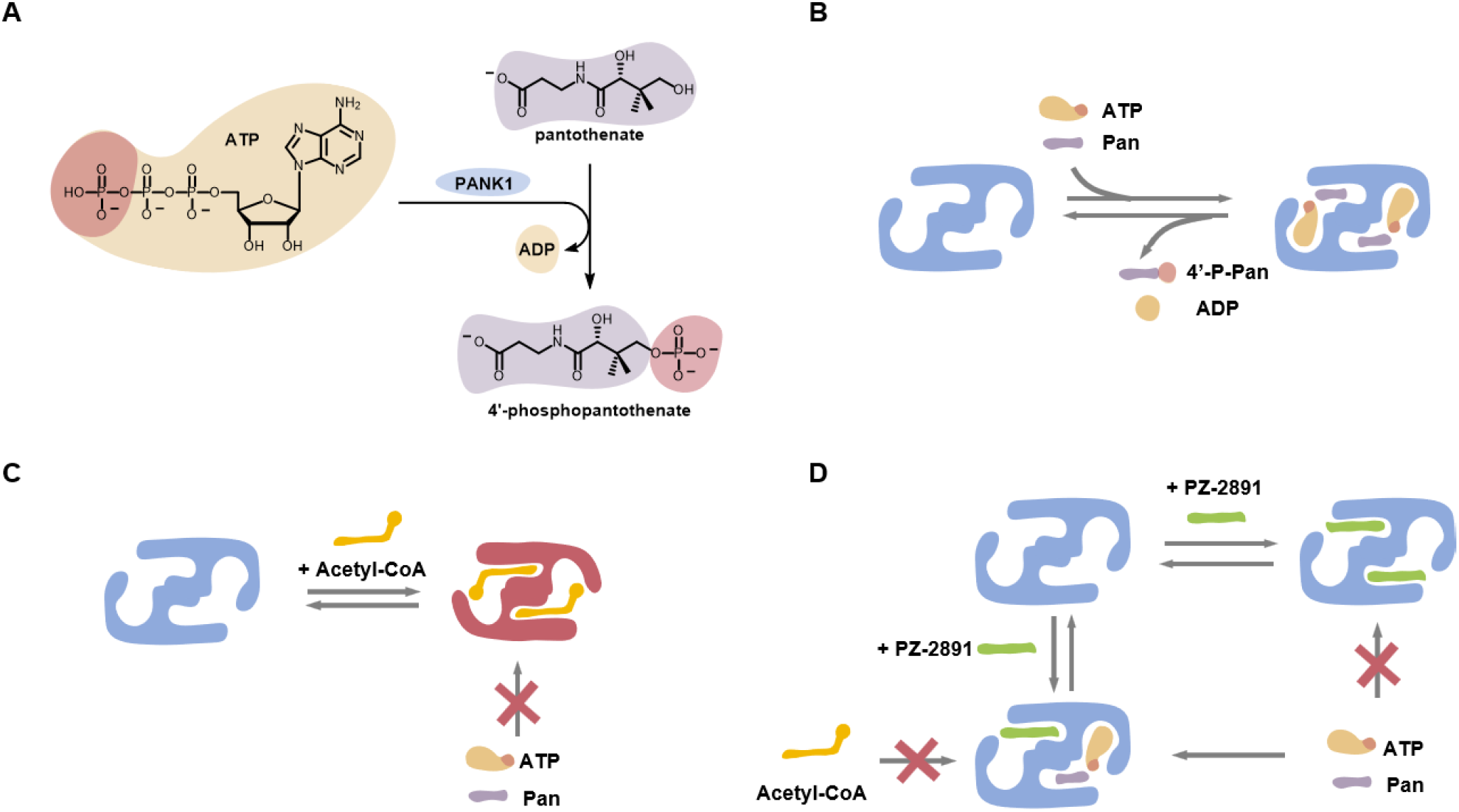
PANK1-catalyzed reaction and its regulation by acetyl-CoA and PZ-2891. (A) PANK1 transfers the terminal phosphate of ATP to the 4’-hydroxyl group of pantothenate, yielding 4’-phosphopantothenate and ADP. (B) Schematic of the PANK1 catalytic cycle showing substrate binding and product release. Pan denotes pantothenate, and 4’-P-Pan denotes 4’-phosphopantothenate. (C) Acetyl-CoA binding leads to the inactivation of PANK1 and inhibition of substrate binding. (D) Proposed model for the dual effects of PZ-2891. Binding to one single site within the PANK1 dimer stabilizes the active conformation, antagonizing acetyl-CoA feedback inhibition while leaving the second site available for catalysis. Occupanying both sites will block substrate access.

## 2. Material and Methods

### 2.1 Constructs

The cDNA encoding human PANK1α (residues 231-598) was cloned into an N-terminal His-tagged pET vector (pPH) to generate a construct carrying N-terminal His and Strep tags.

### 2.2 Expression and purification of PANK1

For protein expression, the pPH-PANK1 construct was transformed into *E. coli* NiCo21(DE3). Cells were cultured in LB medium at 37□ □. When the cell density reached OD_600_□ □= □ □0.7, the culture was cooled to 18□ □for 30□min. After that, 1□mM isopropyl-β-d-thiogalactoside was added and the culture was incubated for an additional 16□h at 18□ □.

Cells were harvested, washed with TBS [20 mM Tris (pH 8.0 at 4□), 150 mM NaCl], and sonicated in lysis buffer containing 50□mM Tris (pH 8.0 at 4 □), 300□mM NaCl, 10□mM MgCl_2_, 1 mM phenylmethylsulfonyl fluoride, and 3.5 nM benzonase. The lysate was clarified by centrifugation at 16,000 rpm for 30 min, and the supernatant was applied to Ni-NTA columns.

The columns were washed sequentially with buffer containing 20 mM Tris (pH 8.0 at 4 □) and 500 mM NaCl, followed by buffer containing 20 mM Tris (pH 8.0 at 4 □), 150 mM NaCl, and 20 mM imidazole. The target protein was eluted with elution buffer containing 150 mM NaCl and 250 mM imidazole (pH 8.0).

The Ni-NTA eluate was subsequently loaded onto Strep-affinity columns pre-equilibrated with TBS supplemented with 5 mM DTT. The columns were washed with the same buffer, and the target protein was eluted with buffer containing 50 mM Tris (pH 8.0 at 4 □), 150 mM NaCl, 5 mM desthiobiotin, and 5 mM DTT. The Strep-column eluate was concentrated and further purified by size-exclusion chromatography using a Superdex 200 Increase column equilibrated with 20 mM Tris (pH 8.0 at 4 □), 200 mM NaCl, and 2 mM DTT. Peak fractions were analyzed by SDS-PAGE, and fractions corresponding to the target protein were pooled. Purified protein was supplemented with glycerol to a final concentration of 20% (v/v) before being flash-frozen in liquid nitrogen and stored at -80 □for subsequent biochemical assays.

### 2.3 Malachite green assay

Ammonium molybdate (Sigma) was dissolved in 4 M HCl to prepare a 4.2% (w/v) solution. Malachite green (Solarbio) was dissolved in water to prepare a 0.045% (w/v) solution. The ammonium molybdate and malachite green solutions were mixed at a volume ratio of 1:3 and stirred for 30 min. Tween 20 was added to a final concentration of 0.1% (v/v). The resulting reagent was filtered through a 0.22-μm membrane and stored at 4 □ for up to one month. Pi standards were prepared from Na_2_HPO_4_ in water at concentrations of 0, 5, 10, 20, and 40 μM. ADP and 4′-phosphopantothenate (4’-P-Pan; TargetMol, Cat. No. T202207-3) solutions were prepared at 40 μM. To each well of a 96-well plate, 50 μl of the sample and 20 μl of water were added. Then 70 μl of Malachite green reagent equilibrated to room temperature was added. After incubation for 30 min at room temperature with shaking at 80 rpm, absorbance at 620 nm was measured using an Infinite 200 Pro microplate reader (Tecan Life Sciences). Water was used as the vehicle control.

### 2.4 ADP-Glo kinase activity assays

PANK1 activity was measured using the ADP-Glo assay kit (Promega; Cat. No. V6930). Reactions were performed in 384-well plates in a total volume of 5 μl containing kinase assay buffer (40 mM Tris, pH 7.5, 20 mM MgCl_2_, and 0.1 mg/mL BSA), 100 μM ATP, 100 μM pantothenate, 50 ng of purified PANK1, and 0.5% (v/v) DMSO, in the presence or absence of 10 μM acetyl-CoA.

Empagliflozin provided by Boehringer Ingelheim, dapagliflozin (TargetMol, Cat. No. T2389-0), and PZ-2891(MCE, Cat. No. HY124634) were dissolved in DMSO and diluted 40-fold in 2× assay buffer to prepare working solutions. Substrate mixtures containing ATP and pantothenate were prepared with or without acetyl-CoA. Reactions were initiated by combining 3 μl of the appropriate substrate mixture, 1 μl of compound solution, and 1 μl of purified PANK1 (50 ng/μl). 0.5% (v/v) DMSO was used as the vehicle control. After incubation of the reaction system at room temperature for 30 min, 5 μl of ADP-Glo Reagent was added to stop the reaction and the mixture was incubated at room temperature for 40 min to deplete residual ATP. 10 μl of Kinase Detection Reagent was added and the mixture was incubated for 30 min to allow luminescence detection of ADP. Luminescence was detected using an Infinite 200 Pro microplate reader (Tecan Life Sciences) with an integration time of 0.5 s. To facilitate comparisons across groups, the non-enzyme background was subtracted from the luminescence signals, which were then normalized to the respective vehicle controls.

## 3. Results

### 3.1 The malachite green assay does not measure PANK1 activity

The recent study used a malachite green assay to measure empagliflozin- and PZ-2891-dependent PANK1 activation^14^. The malachite green assay is a colorimetric method designed to quantify free Pi^17^. Under strong acidic conditions, Pi reacts with molybdate to form a molybdophosphoric anion, which associates with the cationic dye malachite green to produce a blue-green ion-association complex with absorbance at approximately 610-630 nm (Fig. 2A). The absorbance signal is proportional to the Pi concentration^17^. However, the reaction catalyzed by PANK1 does not generate Pi, because its products are ADP and 4′-phosphopantothenate.

**Fig. 2.**
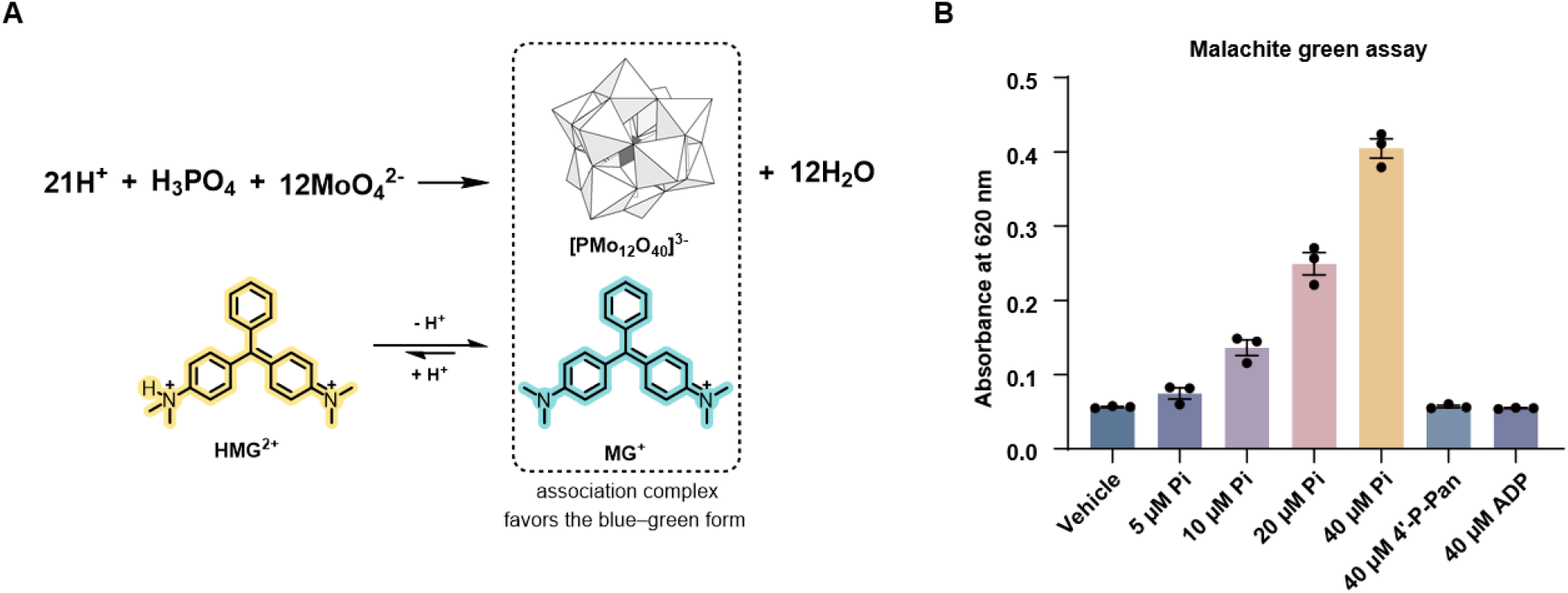
Principle of the malachite green assay. (A) Under acidic conditions, inorganic phosphate reacts with molybdate to form phosphomolybdate ([PMo_12_O_40_]^3-^). Association with phosphomolybdate stablizes the blue-green form of malachite green (MG^+^), which can be quantified by its absorbance at 620 nm. (B) Absorbance at 620 nm in the Malachite green assay with inorganic phosphate (Pi; 5-40 µM), 4’-phosphopantothenate (4’-P-Pan; 40 µM), or ADP (40 µM); vehicle (H_2_O) served as the blank control. Data are presented as mean ± SEM. n = 3 independent experiments.

To verify that the malachite green assay cannot detect either product of the PANK1 reaction, ADP and 4′-phosphopantothenate were individually incubated with the assay reagents. As expected, Pi produced a robust signal in a concentration-dependent manner, whereas neither ADP nor 4′-phosphopantothenate generated a detectable signal above the blank control under the conditions tested (Fig. 2B). These observations indicate that absorbance measured in a malachite green-based PANK1 assay cannot reflect PANK1-catalyzed pantothenate phosphorylation. Any Pi detected under such conditions likely originates from ATP hydrolysis by a contaminating ATPase or from nonenzymatic ATP degradation.

### 3.2 SGLT2i do not activate PANK1 in an ADP-Glo luminescent assay

Because the malachite green assay cannot measure the activity of PANK1, we used the established ADP-Glo Kinase Assay^18^, a luminescent method that quantifies the ADP generated during PANK1-catalyzed reaction. After termination of the enzymatic reaction, residual ATP is depleted, and the detection reagent converts the ADP back into ATP, which is then consumed by a coupled luciferase/luciferin reaction. The resulting luminescence is proportional to the amount of ADP and therefore to PANK1 catalytic activity^18^.

Recombinant human PANK1 was expressed in *E. coli* and purified sequentially by Ni-NTA affinity chromatography, Strep-Tactin affinity chromatography, and size-exclusion chromatography. These sequential chromatographic steps removed several contaminant proteins (Fig. 3A). The final protein preparation exhibits high purity on SDS-PAGE (Fig. 3A) and was used for activity measurements. In the absence of acetyl-CoA, an endogenous inhibitor, purified PANK1 exhibited robust basal activity. Consistent with previous results^16^, PZ-2891 can inhibit this activity, reflecting the competitive inhibition by binding to the substrate binding pocket (Fig. 3B). In contrast, neither empagliflozin nor dapagliflozin altered PANK1 activity at the concentrations tested (Fig. 3B). We next evaluated enzyme activity in the presence of acetyl-CoA. As expected, substoichiometric PZ-2891 relieved acetyl-CoA-mediated feedback inhibition and thus activated PANK1, reproducing the characteristic one-protomer allosteric activation described previously (Fig. 3B)^16^. However, under the same conditions, empagliflozin and dapagliflozin showed no measurable activation of PANK1 (Fig. 3B). Collectively, these results demonstrate that SGLT2i do not activate purified PANK1 either in the basal state or under acyl-CoA feedback inhibition, in great contrast to the direct activation reported in the recent study^14^.

**Fig. 3.**
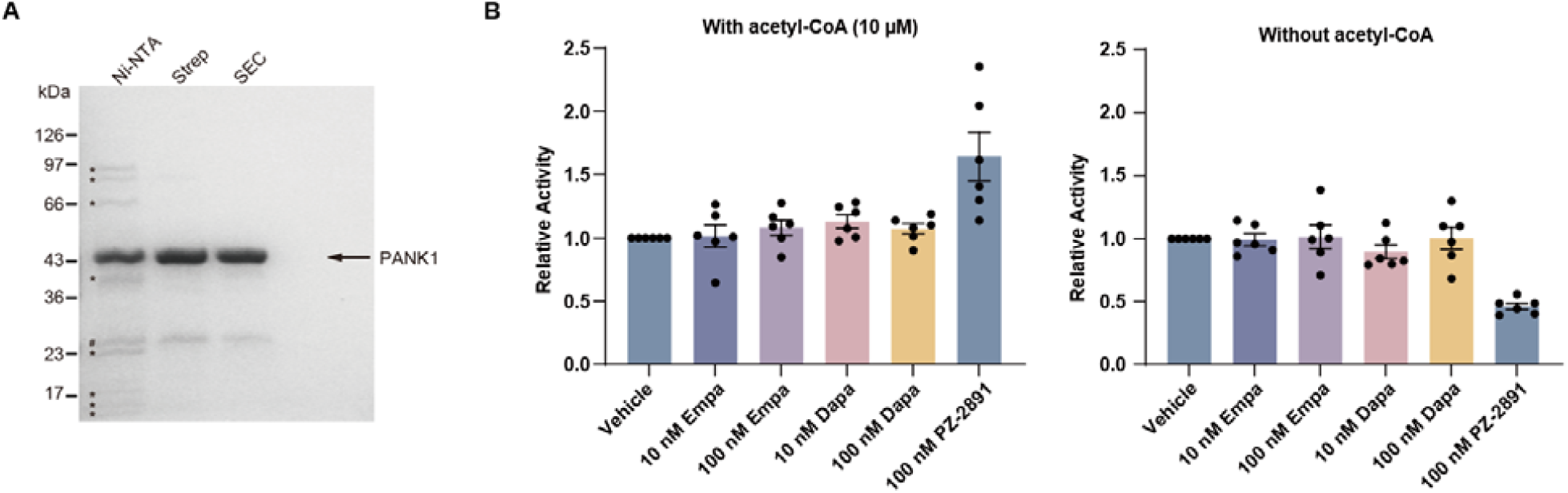
Effects of SGLT2 inhibitors on purified PANK1. (A) SDS–PAGE analysis of recombinant PANK1 after sequential purification by nickel-affinity chromatography (Ni-NTA), Strep-tag affinity chromatography (Strep), and size-exclusion chromatography on a Superdex 200 column (SEC). 3 µg eluates from each purification step were loaded onto SDS-PAGE. Asterisks denote contaminating proteins. The hash symbol indicates a putative partial degradation fragment of PANK1. Molecular masses are indicated in kDa. (B) PANK1 activity measured using the ADP-Glo assay in the absence or presence of acetyl-CoA (10 µM) after treatment with empagliflozin (Empa; 10 or 100 nM), dapagliflozin (Dapa; 10 or 100 nM), or PZ-2891 (100 nM). To facilitate comparisons across groups, all signals were corrected for the corresponding blank background and normalized to their respective vehicle controls (0.5% (v/v) DMSO). Data are presented as mean ± SEM. n = 6 independent experiments.

## 4. Discussion

In addition to the biochemical assays of purified PANK1, the recent study reported that empagliflozin promoted CoA synthesis in perfused human cardiac tissue and isolated cardiomyocytes through direct activation of PANK1^14^. Our biochemical reanalysis does not reproduce a direct activation effect of SGLT2i on PANK1. In the malachite green assay used in that report^14^, the assay measures free Pi and signals attributed to PANK1 activation therefore likely reflect ATP hydrolysis by a contaminating ATPase rather than pantothenate phosphorylation. When PANK1 activity was monitored by detecting its stoichiometric ADP product using the ADP-Glo assay, SGLT2i failed to activate purified PANK1 under either basal or acetyl-CoA-inhibited conditions.

Our findings do not rule out the possibility that SGLT2i modulate cardiac CoA metabolism in intact tissue. Such effects, if present, could arise through mechanisms unrelated to direct PANK1 activation, for example altered pantothenate uptake or availability, regulation of other CoA biosynthetic enzymes, activation of upstream signaling pathways that control CoA homeostasis, changes in substrate supply. The cardiac metabolic phenotypes reported in tissue and cellular systems should therefore be interpreted in light of the present observation that PANK1 itself is not a direct target of SGLT2i.

In summary, our findings show that SGLT2i do not act as direct activators of PANK1 under the conditions tested. The previously proposed PANK1-dependent mechanism accounting for the cardiovascular benefits of SGLT2 inhibitors therefore requires reconsideration, while the molecular basis of SGLT2i cardiovascular benefits remains to be elucidated.

## Credit authorship contribution statement

Zihao Guo: Methodology, Validation, Formal analysis, Investigation, Visualization, Writing - review & editing; Jiahe Zang: Methodology, Validation, Formal analysis, Investigation, Writing - review & editing; Lei Chen: Conceptualization, Methodology, Validation, Formal analysis, Investigation, Writing original draft, Writing - review & editing, Visualization, Supervision, Project administration, Funding acquisition.

## Conflict of Interest

All authors have no conflicts of interest.

## Acknowledgement

We thank Stefan Hoerer and Herbert Nar from Boehringer Ingelheim for providing the empagliflozin.

## Funding

The work is supported by grants from the Ministry of Science and Technology of China (National Key R&D Program of China, 2022YFA0806504 to L.C.), the National Natural Science Foundation of China (32430050, 32225027 to L.C.), Center For Life Sciences (CLS to L.C.) and The Fundamental Research Funds for the Central Universities, Peking University to L.C.

## Declaration of generative AI use

During the preparation of this manuscript, the authors used Doubao to improve the language, grammar, and readability of the manuscript. The authors reviewed and edited all AI-assisted output and take full responsibility for the accuracy, originality, and integrity of the final manuscript.

## Abbreviations

SGLT2: Sodium-glucose cotransporter 2
SGLT2i: SGLT2 inhibitors
PANK1: pantothenate kinase 1
CoA: coenzyme A
T2DM: type 2 diabetes mellitus
HFrEF: heart failure with reduced ejection fraction
HFpEF: heart failure with preserved ejection fraction
Pi: inorganic phosphate
DTT: dithiothreitol.

## Notes

### Competing Interest Statement

The authors have declared no competing interest.

